# CD4 Co-Receptor Regulates Sex-Specific NK Cell Responses to Acute *Toxoplasma gondii* Infection

**DOI:** 10.1101/2024.12.06.627254

**Authors:** Tathagato Roy, Leah Bernstein, Hunter K. Keplinger, Kaatje Fisk, Sai K. Ng, Stephen L. Denton, Jason P. Gigley

**Affiliations:** Department of Molecular Biology, University of Wyoming, Laramie, WY, U.S.A

**Keywords:** NK cells, Sexual Dimorphism, *T. gondii*

## Abstract

Immunity to *Toxoplasma gondii* (*T. gondii*) is sexually dimorphic in humans and mice, with females having higher morbidity and mortality during immune dysfunction and HIV-AIDS. The mechanisms underlying these sex differences are unclear. We investigated how a lack of CD4+ T cells (CD4 co-receptor KO) impacted *T. gondii* survival in mice. Female CD4 co-receptor KO mice succumbed to *T. gondii* much faster than males. To dissect why female CD4 co-receptor KO mice died faster, we tested their NK cell responses to acute *T. gondii* infection compared to males. Although in wild-type (WT) animals, both sexes had similar increases in total NK cells and IFNγ + NK cells, infected CD4 co-receptor KO female mice had 50% fewer IFNγ+ NK cells than infected WT female mice. Infected male CD4 co-receptor KO had a similar increase in IFNγ+ NK cells as WT male mice. Since CD4 co-receptor deficient mice still have functional helper T cells that are CD4−, we next tested survival and NK cell responses in female and male MHCII deficient (MHCIIKO) animals, which completely lack helper CD4+T cells. Surprisingly, survival, NK cell numbers, and IFNγ+ NK cells were not significantly different between WT or MHCIIKO female and male mice. These results suggest CD4 co-receptor expression is required for survival via optimal NK cell responses during acute *T. gondii* infection only in female mice and not in male mice. Our findings reveal an unappreciated sexual dimorphic role of CD4 co-receptor expression in regulating NK cell responses to acute *T. gondii* infection.

## Introduction

*Toxoplasma gondii* (*T. gondii*) is an obligate intracellular parasite that infects approximately 30% of humans [1]. *T. gondii* infection is a major cause of foodborne illness resulting in hospitalization; however, after the acute stage of infection, most healthy individuals are asymptomatic. The parasite is a severe health threat to immune-compromised patients such as HIV/AIDS, organ transplantation, cancer, or those who have received immune-suppressive therapies [2–6]. Uncontrolled acute infection and reactivated chronic infection in these patients can result in Toxoplasmic Encephalitis (TE) and death. *T. gondii* is also a significant health threat during pregnancy. The parasite can pass the blood-placental barrier to infect the developing fetus, resulting in congenital toxoplasmosis [7]. Congenital toxoplasmosis can lead to visual impairment, mental disability, and fetal loss through embryonic death or spontaneous abortion [7–11]. Currently, there are no fully effective treatments or vaccines that can completely prevent and clear this infection, and once infected, the infection persists throughout the host’s life [12]. A robust innate immune response is critical to limit chronic *T. gondii* infection, especially in immune-compromised individuals [1, 13].

Immunity develops against *T. gondii* infection after TLR recognition of the parasite by neutrophils, macrophages, and dendritic cells (DC) [14–19] [16, 20–22]. This recognition results in the production of the proinflammatory cytokine IL-12, which then activates Natural killer (NK) cells to produce IFNγ. Via their IFNγ production, they act as the first line of innate immune defense against acute *T. gondii* infection to limit parasite replication and dissemination [23, 24]. They have also been shown to provide secondary protection dependent upon IL-12 and IL-23 during lethal parasite challenge in a vaccine challenge model [25]. NK cells are a part of the group 1 innate lymphoid cells (ILCs), which share some phenotypic and functional characteristics with ILC1s. ILC1 have also been implicated to play a role in early control of *T. gondii* infection [26, 27]. Despite this knowledge, how other factors regulate the activation of NK cells during acute *T. gondii* infection is still unexplored.

NK cells are very plastic in their responsiveness and have a complex role in the development and maintenance of immunity [28]. One population of immune cells that they appear to have a particularly interesting relationship with are CD4+ T cells. NK cells can act as surrogate helper cells in the development of CD8+ T cell responses to *T. gondii*. In the absence of CD4+ T cells, NK cells via their IFNγ help prime CD8+ T cells during parasite infection [29]. NK cells also can be negative regulators of CD4+ T cells during persistent infections [30–36]. On the flip side, previous studies have demonstrated that CD4+ T cells can both positively and negatively regulate NK cell responses [37–43]. NK cell function and numbers are enhanced by IL-2 producing CD4+ T cells [37–39]. Infection with *Plasmodium falciparum*, *Leishmania major*, and *Pneumocystis murina* indicate that CD4+ T cells indirectly affect NK cell activation via CD4 co-receptor-MHCII interactions on DCs, resulting in higher DC maturation and IL-12 production [39–42]. Lastly, regulatory CD4+ T cells can also regulate NK cell responses by utilizing IL-2 and limiting this cytokine availability for NK cells to use [43, 44]. There are still many unanswered questions about how CD4+ T cells impact the development of NK cell responses during acute infection.

Although there is extensive knowledge of the IL-12 - NK cell IFNγ axis during infection, most of this information was discovered using only female mice. More recent discoveries have revealed that *T. gondii* infection is sexually dimorphic. Female and male mice have different levels of cytokine production and dissimilar morbidity and mortality outcomes [45–47]. In humans, female HIV-AIDS patients are more susceptible to *T. gondii* infection that causes TE [48, 49]. HIV+ patients progress from having CD4+ T cells to developing AIDS, where their CD4+ T cell counts are very low [50]. This raises the possibility that in females, CD4+ T cells may be more critical for optimal NK cell responses during acute infections than in males. Currently, there is little understanding of the sexual dimorphisms in immunity to *T. gondii* infection and how they contribute to different infection outcomes based on sex. Moreover, the sex-specific role of CD4+ T cells modulating NK cell responses during acute *T. gondii* infection remains unclear.

Our study addresses questions about sexual dimorphism in NK cell dependent immunity to *T. gondii* and how CD4+ T cells play a role in this process. NK cells are plastic in their responsiveness and have a complex role in the development and maintenance of immunity to *T. gondii* infection [27, 28, 51]. In this study, we discovered that CD4 co-receptor expression, but not CD4+ T cells or MHCII, is essential for NK cell response. This intrinsic need for CD4 co-receptor expression for optimal NK cell IFNγ production is observed only in female mice and not in male mice. Our findings reveal an unappreciated sexual dimorphic role of CD4 co-receptor expression in regulating NK cell responses to acute *T. gondii* infection. Consistent with previous literature, our study also demonstrates that females are better at controlling parasite growth and replication during acute infection [46]. Our study is not only important for understanding NK cell responses in immune-compromised patients, especially HIV-AIDS female patients, but also reveals an intrinsic requirement of CD4 co-receptor for optimal NK cell responses in female mice during acute *T. gondii* infection.

## Materials and Methods

### Ethics Statement

All the animal husbandry was carried out by Gigley Laboratory at the University of Wyoming, following Animal Care and Use Committee (IACUC)-approved protocols. To ensure compliance with the U.S. Department of Health and Human Services Guide for the Care and Use of Laboratory Animals, the protocols were approved by the University of Wyoming IACUC (Public Health Service/National Institutes of Health/Office of Laboratory Animal Welfare Assurance number: D16-00135). The Institutional Review Committee granted ethical approval following the eighth edition of the NIH Guide for the Care and Use of Laboratory Animals. An attempt was made to reduce the pain and distress of these animals. High ethical standards were maintained and overseen by the University of Wyoming’s IACUC and members of the Gigley lab.

### Mouse Strains

The mouse strains used in this study included WT (C57BL/6J, #000664), RAG1KO (B6.129S7-Rag1tm1Mom/J, #002216), CD4 co-receptor KO (B6.129S2-Cd4tm1Mak/J, #002663), MHCIIKO (B6.129S2-H2dlAb1-Ea/J, #003584), CBA (CBA/J, #000656), Eomes fl (B6.129S1(Cg)-*Eomes^tm1.1Bflu^*/J, # 017293), Tbx21^F^ (B6.129-*Tbx21^tm2Srnr^*/J, #022741) all purchased from The Jackson Laboratory. NKp46iCre mice were generously provided by Dr. Eric Vivier from the Centre d’Immunologie de Marseille-Luminy, Aix Marseille Université, Hôpitaux Universitaires de Marseille, Marseille, France. NKp46iCre mice were bred with Eomes ^f^/^f^ and T-bet ^f^/^f^ mice to generate NKp46iCre^+/−^Eomes ^f^/^f^ and NKp46iCre^+/−^T-bet ^f^/^f^ mice. Mice were genotyped by PCR after genomic DNA was processed using DirectPCR Digestion Buffer and Proteinase K (supplied by Viagen Biotech). The following primer pairs were used: NKp46iCre Forward (0.2µM): 5’- AGA TGC CAG GAC ATC AGG AAC CTG-3’, NKp46iCre Reverse (0.2µM): 5’- ATC AGC CAC ACC AGA CAC AGA GAT C -3’. The presence of Eomes ^f^/^f^ and T-bet ^f^/^f^ alleles was confirmed using the following primers: Eomes fl Forward (0.3µM): 5’- AGA CAG TTT TTG GAG CCA GAT G-3’, Eomes fl Reverse (0.3µM): 5’- CCT CTT TGG GTC CCT GTC TC-3’, and T-bet fl Forward (0.5µM): 5’- AGT CCC CCT GGA AGA ACA CT-3’, T-bet fl Reverse (0.5µM): 5’- TGA AGG ACA GGA ATG GGA AC-3’. NKp46iCre^−/−^ Eomes ^f/f^ and NKp46iCre^−/−^T-bet ^f/f^ littermates were used as controls, and 8–12-week-old mice were used for experiments.

### T. gondii strains and Infection

Type I *T. gondii* RH strain was sustained through regular passages in MRC5 fibroblasts cultured in 1% Dulbecco’s Modified Eagle Medium (1% DMEM) supplemented with 1% Fetal Bovine Serum, 1X Penicillin/Streptomycin, and 1X Amphotericin B. After lysis of the MRC5 monolayer, tachyzoites were purified by 3-micron filtration and counted using a hemocytometer. Type II strain ME49 was maintained by serial passage in 8-week-old CBA mice. Five weeks post-infection, the infected CBA mouse brain was homogenized using a Dounce homogenizer, and cysts were enumerated on a microscope. All infections were administered in 200 μL sterile 1X phosphate buffered saline (PBS). To perform survival studies, CD4KO, RAG1KO, and MHCIIKO mice were infected intraperitoneally (i.p.) with 2-3 cysts of ME49 strain, and for all other acute studies, mice were injected (i.p.) with 10-20 cysts ME49 (5-week-old). A clinical scoring system was utilized to assess the health status of the mice post-infection. Mice were assigned a numerical score based on observed clinical symptoms as follows: 0 indicated normal behavior; 1 represented mice with slightly ruffled fur; 2 was assigned to mice displaying a swollen abdomen and ruffled fur; 3 denoted mice showing an arched back, swollen abdomen, and ruffled fur; 4 indicated mice with slightly reduced activity, along with the aforementioned symptoms; and 5 was designated for mice exhibiting low activity, labored breathing, closed eyes, an arched back, swollen abdomen, and ruffled fur. Mice scoring above 4 were promptly euthanized to ameliorate further suffering and distress. The weights of mice were measured every other day after infection.

### Parasite Burden by Real-Time PCR

Peritoneal exudate cells (PEC) were harvested by washing the peritoneal with 7 mL of ice cold sterile 1 X PBS. PEC DNA was extracted following the manufacturer’s instructions on days five and twelve post-infection using the DNeasy Blood & Tissue Kit (Qiagen 69506). The concentration and quality of the extracted DNA were quantified using a Nanodrop Spectrophotometer (NanoVueTM Plus Spectrophotometer, VWR). Parasite burdens were quantified in 400 ng of total DNA extracted from PEC per sample and run in triplicate. Parasite gene B1 was amplified employing SSO Advanced SYBR Master Mix (Bio-Rad 1725017) with 0.5μM of Forward (5’- CGT CCG TCG TAA TAT CAG -3’) and Reverse (5’- GAC TTC ATG GGA CGA TAT G -3’) primers as previously published [52].

### ELISA

Eye bleeding was performed to collect the blood samples from experimental animals under isoflurane-induced anesthesia (VetOne 502017). The blood was coagulated within 30 minutes at room temperature (RT). The blood was centrifuged to separate the serum and used for assay. PEC and Spleen cells isolated and resuspended in complete Iscoves DMEM (10% FBS, 1 X Penn/strep, 1X Amphotericin B, L-glutamine, non-essential amino acids, sodium pyruvate, and 2-mercaptoethanol) media at 2×10^6^ PEC cells/well or 5×10^6^ spleen cells/well. Cells were incubated for 24 hours at 37°C with 5% CO_2_. Cells were lysed by repeated freeze-thaw 3 times, then cells were pelleted by centrifugation, and the supernatant was collected. Spleen and PEC samples were diluted to 1:20 and 1:40 and analyzed for IL-12p40 by ELISA (BioLegend 431604) per the manufacturer’s instructions.

### Single-cell Suspension and Flow Cytometry

PECs were isolated and resuspended in 1 X Stain/Wash Buffer with EDTA (SWBE, containing 2% w/v Bovine Calf Serum and 2 mM EDTA in 1 X PBS). Spleen cells were isolated, then resuspended in 1 mL of 1X RBC Lysis Buffer (Tonbo Biosciences #TNB-4300-L100) for 3 minutes with gentle agitation to lyse the red blood cells (RBC) and quenched with 1X SWBE. PEC and splenocytes were resuspended at 1 X 10^7^ per ml, and 1 X 10^6^ cells were plated per well for staining procedures. Cells were initially stained with fixable Live/Dead Aqua viability dye (Invitrogen) following the manufacturer’s instructions. Cells were then surface stained with diluted antibodies in 1X SWBE with Anti-CD16/32 FcR blockade to reduce nonspecific staining. Cells were then fixed using the BD Cytofix/Cytoperm™ Fixation/Permeabilization kit according to the manufacturer’s recommendations. Cells were either analyzed by flow cytometry for phenotype or were further stained intracellularly after permeabilization for functional analyses. NK functional assays were performed in plates pre-coated with 0.5μg per well of NK1.1 (PK136, BioXCell BE0036). Cells were stimulated for 4 hours at 37°C and 5% CO_2_ for 4 hours with 0.66X Protein Transport Inhibitor Cocktail (PTIC, eBioscience 00-4980-03), 1X Cell stimulation cocktail (eBioscience 00-4970-93) in complete Iscove’s DMEM with L-Glutamine and 25mM HEPES supplemented with 1X Penicillin/Streptomycin, Amphotericin B, Non-Essential Amino Acids, 0.1mM Sodium Pyruvate, 0.2mM Glutamine XL, 10% Fetal Bovine Serum, and 0.1mM 2-mercaptoethanol. Cells were then surface and intracellularly stained and analyzed by flow cytometry. The following extracellular staining were used: CD3-FITC (17A2, BioLegend 100204), CD335/NKp46-APC (29A1.4, BioLegend 137608), and CD49b-Biotin (DX5, BioLegend 108904). The secondary and intercellular stains were Streptavidin-PE/Cy7 (BioLegend 405206) and IFNg-PE (XMG1.2, BioLegend 505806), respectively. Cells were run on a Guava 12 HT flow cytometer (Cytek) and analyzed using Flowjo.

### Statistical Analysis

Statistical data analysis was performed using Microsoft Excel 2023, Prism 6.0h, and Prism v.10.1.0(264) (GraphPad, La Jolla, CA). Survival rates were quantified utilizing the Log-rank (Mantel-Cox) test. For datasets with three or more normally distributed groups, ordinary one-way ANOVA (with nonparametric or mixed methods) was applied, followed by Turkey’s multiple comparisons test. To compare the normal distribution data sets with unequal variances, a student’s t-test with Welch’s correction was performed. The mean with the standard deviation (SD) was graphically denoted to better visually represent the data sets. “ns” was not significant (p > 0.05). In contrast, significance was indicated by a maximum p-value of 0.05 or below. A confidence interval of 95% acted as the cutoff to identify the significant differences between groups: *p ≤ 0.05, **p ≤ 0.01, ***p ≤ 0.001, ****p ≤ 0.0001.

## Results

### CD4 Co receptor is required for optimal NK cell response in female mice but not in males during acute T. gondii infection

Previous studies have shown that *T. gondii* infection is sexually dimorphic [45–47]. Female HIV-AIDS patients are more prone to *Toxoplasma* infection [48]. In *T. gondii* infection, IFNγ+ Natural Killer (NK) cells are the major early controller of acute infection [23, 24]. Previous studies demonstrate that CD4+T cells can directly or indirectly regulate NK cell activity [39–42]. This raises the possibility that the reason why female patients are more susceptible to *T. gondii* infection during HIV/AIDS could be due to defective NK cell responses. Additionally, how CD4+T cells regulate NK cell activity during the acute phase of *T. gondii* infection is not known. To test these questions, WT and CD4 co-receptor KO females and males were infected intraperitoneally with 20 cysts of *T. gondii* (type II strain of the parasite, ME49). On day five post-infection, the percentage and absolute number of NK cells (Lin-CD49b+NKp46+) and IFNγ+ NK cells were measured at the site of infection (peritoneum) by flow cytometry. NK cells increased in all infection groups; however, the absence of the CD4 co-receptor resulted in a nearly 50 % decrease in the percentage and absolute number of NK cells in the peritoneum upon infection in female mice (Figure 1A and B) compared to male mice. Additionally, CD4 co-receptor KO female mice had a lower fold increase in the percentage and absolute number of NK cells after infection compared to WT female mice at the peritoneum (Figure 1C). Since IFNγ is the major mediator of *T. gondii* infection [53], we measured the percentage and absolute number of IFNγ+ NK cells at the site of infection. Upon infection, the percentage (Figure 1A and D) and absolute number (Figure 1D) of IFNγ+ NK cells at the peritoneum were higher in both infected female groups. The absence of the CD4 co-receptor did not impact the percentage of IFNγ+ NK cells in female mice; however, there was an approximate 50% lower absolute number of IFNγ+NK cells in the peritoneum during *T. gondii* infection. Additionally, CD4 co-receptor KO female mice had a higher fold increase in the percentage, but not in absolute number, of IFNγ+NK cells after infection compared to WT female mice at the peritoneum (Figure 1E).

**Figure 1.**
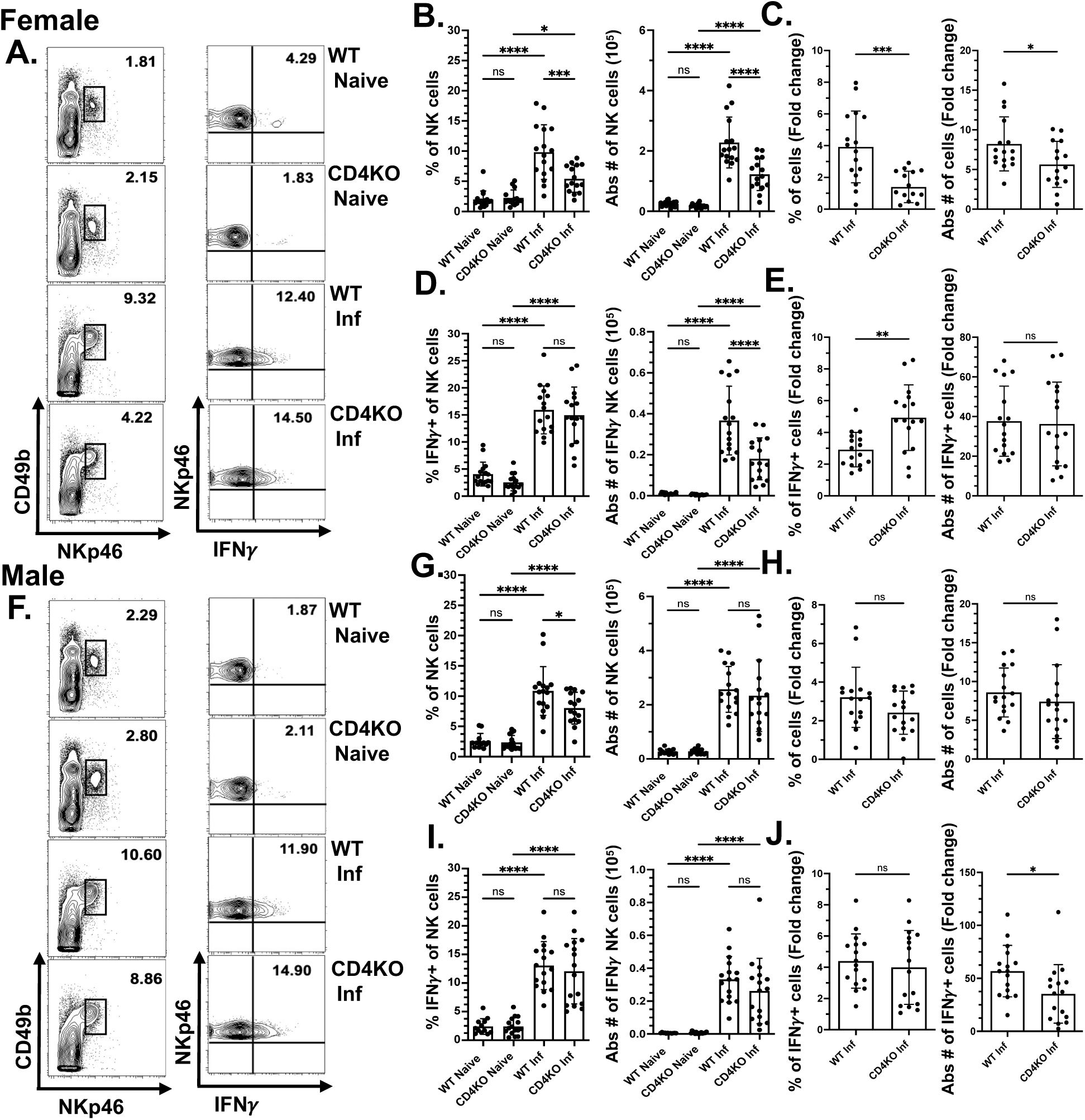
CD4 Co-receptor is required for optimal NK cell response in female mice but not in males during acute T. gondii infection. WT and CD4KO females and males were infected intraperitoneally with 20 cysts of *T. gondii* (type II strain of the parasite, ME49). Five days post-infection, peritoneal cells were harvested and stained for Live/Dead, CD3, NKp46, CD49b, and IFN*γ* to identify NK cells using flow cytometry. NK cells (Lin-NKp46+CD49b+) and IFN*γ*+NK cells (Lin-NKp46+CD49b+IFN*γ*+) were measured and quantified at the infection site in each group. Representative contour plots of peritoneal NK cells and IFN*γ*+NK cells are shown: **(A.)** females, **(F.)** males. Graphs represent **(B.)** the percentage and absolute number of NK cells in females and **(G.)** the percentage and absolute number of NK cells in males in non-infected versus infected WT and CD4KO mice. Fold change increase of percentage and absolute number of NK cells upon infection are shown in **(C.)** females and **(H.)** males. Similarly **(D.)** the percentage and absolute number of IFN*γ*+NK cells in females and **(I.)** the percentage and absolute number of IFN*γ*+NK cells in males in non-infected versus infected WT and CD4KO mice are presented in the graphs. Fold change increase of percentage and absolute number of IFN*γ*+NK cells upon infection are shown in **(E.)** females and **(J.)** males. Data were pooled from 4 individual experiments. Ordinary one-way ANOVA was used to evaluate the NK cell populations. The data are presented as mean ± SD. *p < 0.05, **p < 0.01, ***p < 0.001, ****p < 0.0001.

Like female mice on day five post-infection, the percentage and absolute number of male NK cells at the site of infection were higher in both infected groups. The absence of the CD4 co-receptor resulted in a slightly lower percentage but not absolute number of NK cells at the peritoneum upon infection in male mice (Figure 1F and G). Also, there was no significant difference in the fold change in percentage or absolute number of NK cells after infection in WT males (Figure 1H) compared to CD4 co-receptor KO male mice. Like females, the percentage and absolute number (Figure 1F and I) of IFNγ+ NK cells at the peritoneum were higher in both infected male groups compared to non-infected controls. The absence of the CD4 co-receptor did not impact the percentage or absolute number of IFNγ+ NK cells in the peritoneum of infected male mice. This was different from female mice (Figure 1F and >I). CD4 co-receptor KO male mice had a higher fold increase in the absolute number, but not in percentage, of IFNγ+NK cells after infection compared to WT male mice at the peritoneum (Figure 1J). WT female and male NK cell responses were compared, and as shown in supplemental figure 1 A-F, there were no significant differences in frequency or absolute numbers of NK cells or IFNγ+NK cells after *T. gondii* infection. These findings indicate that the CD4 co-receptor is required for an optimal NK cell response, which is sexually dimorphic during acute *T. gondii* infection.

### CD4 co-receptor Knockout females are more prone to T. gondii infection

To test whether the absence of the CD4 co-receptor results in differential sex-specific survival outcomes in *T. gondii* infection, mice were infected orally with 2-3 cysts of ME49. A physiologically relevant low dose of 2-3 cysts was used instead of the typical 10-20 cysts because it elicits a better long-term immune response in *T. gondii* infection [54]. Without the CD4 co-receptor, females were more susceptible to low dosages of *T. gondii* infection. Surprisingly, all female mice died within day thirty-seven and males fifty-four post-infection (Figure 2A). Despite their survival difference, the two groups developed illness (depicted as sickness score) to infection in a similar fashion (Figure 2B). While aged-matched male CD4 co-receptor KO mice had higher weight than females (till 34-day post-infection/DPI) (Figure 2C), on day 18-30 and 34-36 post-infection, females lost weight more dramatically than CD4 co-receptor KO infected males (Figure 2D). We then measured how the absence of CD4 co-receptors in female and male mice impacted acute parasite burdens. The absence of the CD4 co-receptor did not correlate with a higher parasite burden in the peritoneum during the acute phase of infection (5 DPI). Additionally, female mice controlled the acute infection better than male mice when the immune system was intact (Figure 2E). This result is consistent with recent findings in BALB/c mice infected intraperitoneally with type II parasites [46].

**Figure 2.**
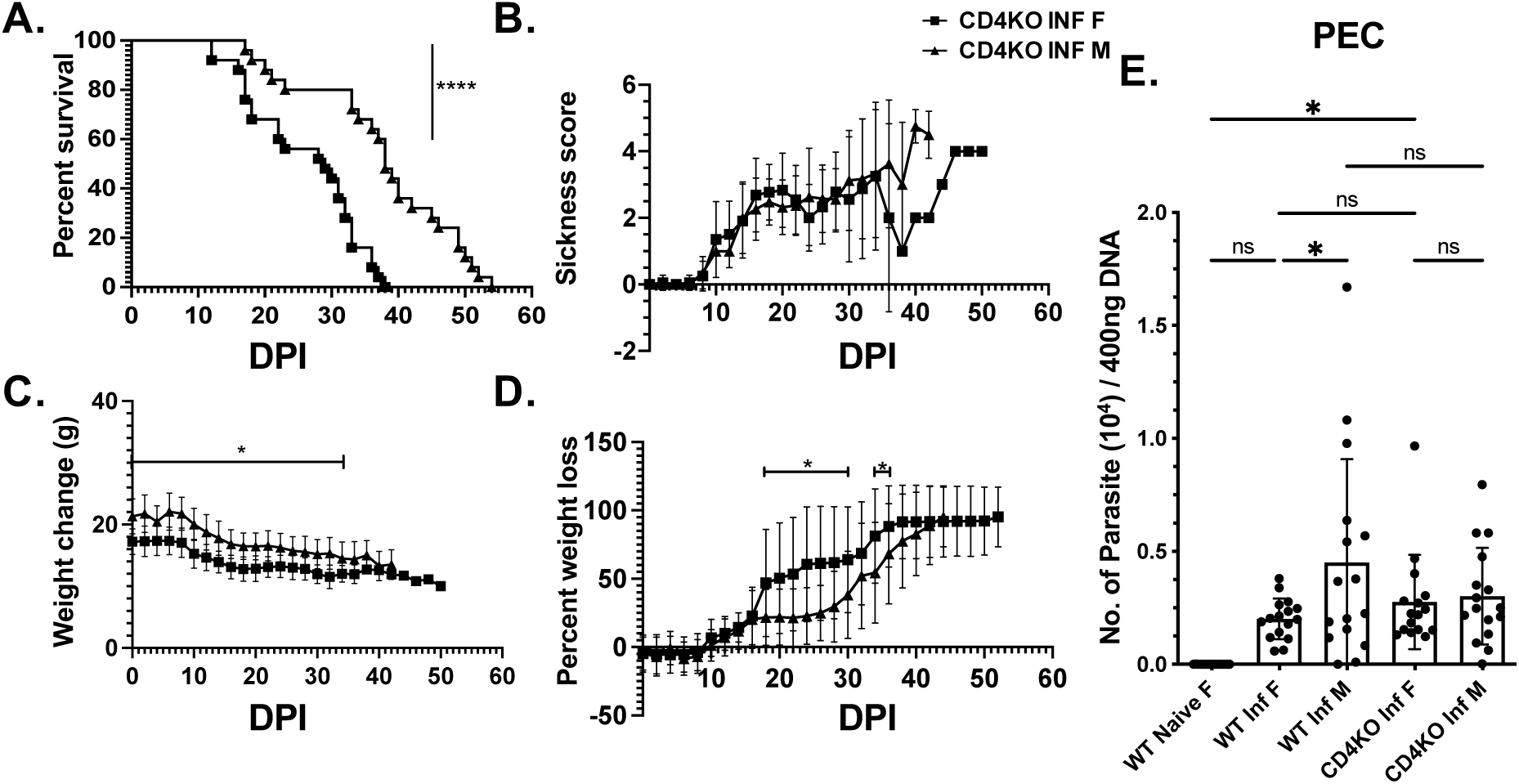
CD4 co-receptor Knockout females are more prone to T. gondii infection. CD4KO female and male mice were infected orally with 2-3 cysts of *T. gondii* (type II strain of the parasite, ME49). Graphs present **(A.)** percent survival of CD4KO females and males (n=25), **(B.)** sickness score, **(C.)** weight change, and **(D.)** weight loss (n=20) measured every other day, till day 50 post-infection. **(E.)** Graph presents *T. gondii* parasite burden of WT and CD4KO female versus male mice on day five post-infection (infection: intraperitoneal with 20 cysts of ME49, mice n=16) in peritoneum. The log-rank (manel-Cox) test was used to evaluate survival rates. Ordinary one-way ANOVA (Brown-Forsythe and Welch) was used to evaluate the parasite burden. Data are ± SD. *p<0.05, **p<0.01, ***p<0.001, ****p<0.0001.

### CD4+T cells are not required for optimal NK cell response in female mice during acute T. gondii infection

Our data demonstrates CD4 co-receptor expression is required for optimal NK cell responses in female but not in male mice during acute *T. gondii* infection. CD4 co-receptor KO mice still have a functional population of double negative CD4− CD8-αβ+ T cells which can still provide help to develop adaptive immunity [55]. To investigate whether CD4+T cells themselves are required for optimal NK cell response in female mice, we repeated our experiments with MHCIIKO female mice. MHCIIKO mice lack all CD4+ T cells in the periphery [56]. WT and MHCIIKO mice were infected intraperitoneally with 20 cysts of ME49, and their NK cell responses were measured at the sight of infection on day 5 post-infection. Our results indicate that in the absence of CD4+ T cells, NK cell percentages and absolute numbers still increased and were comparable to NK cell responses in WT control animals (Figure 3A and C). This was the same for IFNγ+NK cells (Figure 3B and D) in the peritoneum in female mice. To test whether the absence of the CD4+ T cells resulted in differential sex-specific survival outcomes in *T. gondii* infection, MHCIIKO female and male mice were infected orally with 2-3 cysts of ME49. As expected, the absence of CD4+T cells does not correlate with sex-specific differential survival outcomes. Both the groups die within 50 DPI (Figure 3E). Although MHCIIKO-infected male mice were heavier than infected females until day 25 post-infection (data not shown), their percent weight loss (Figure 3F) was identical and followed the same pattern. Just before day 10 post-infection, both groups started losing weight until day 14, after which they recovered before experiencing a progressive and steady weight loss from 20 DPI. Both groups developed illness to infection similarly (Figure 3G), consistent with their weight loss.

**Figure 3.**
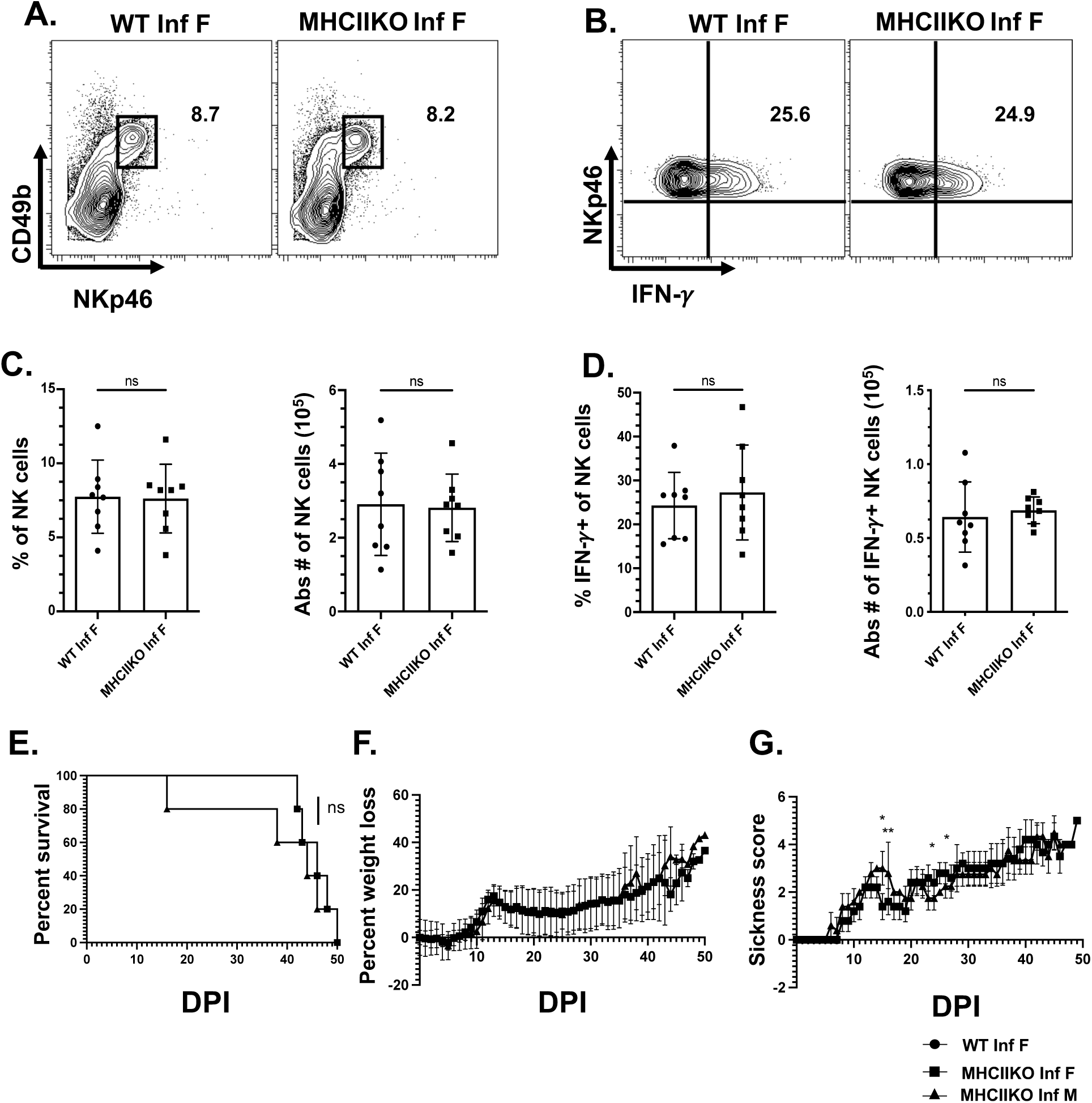
CD4+T cells are not required for optimal NK cell response in female mice during acute T. gondii infection. WT and MHCIIKO female mice were infected interperitoneally with 20 cysts of *T. gondii* (type II strain, ME49). Five days post-infection, the peritoneal cells were harvested and stained for Live/Dead, CD3, NKp46, CD49b, and IFN*γ*. NK cells were identified using flow cytometry. Female NK cells (Lin-NKp46+CD49b+) and IFN*γ*+NK cells (Lin-NKp46+CD49b+IFN*γ*+) were measured at the site of infection in each group. Representative contour plots of **(A.)** peritoneal NK cells and **(B.)** peritoneal IFN*γ*+ NK cells. Graphs present the percentage and the absolute number of **(C.)** NK cells in the peritoneum and percentage and the absolute number of (**D.)** IFN*γ*+NK cells in the peritoneum. Data pooled from 2 individual experiments. MHCIIKO female and male mice were infected with 2-3 cysts of *T. gondii* (type II strain, ME49). Graphs present **(E.)** the percent survival of MHCIIKO female and male mice (n=5), **(F.)** percent weight loss, and **(G.)** sickness score (n=10). The log-rank (mantel-Cox) test was used to evaluate survival rates. Ordinary one-way ANOVA was used to evaluate the NK cell population. The data are ± SD *p< 0.05, **p<0.01, ***p<0.001, ****p<0.0001.

### Mature lymphocytes are not involved in sexual dimorphic NK cell response during acute T. gondii infection

Previously using MHCIIKO mice, we showed that CD4+T cells are not involved in sexually dimorphic NK cell responses during acute *T. gondii* infection. However, MHCIIKO mice have functional CD8+T and B cells [56], and NK cells could also be impacted by these cells. NK cells do interact with these cells and have been shown to negatively regulate CD8+ T cells to promote immune exhaustion during the chronic phase of *T. gondii* infection [57]. Moreover, B cells and NK cells can cross regulate each other in several different infections [58–67]. In addition, previous studies suggest that B cells can stimulate the IFNγ production of NK cells in vitro [66]. Human NK cells can stimulate autologous resting B cells to produce immunoglobulins, including the switching to IgG and IgA [67]. To test whether mature lymphocytes are involved in sexual dimorphic NK cell responses during acute *T. gondii* infection, we measured NK cell responses in parasite-infected RAG1KO mice that lack mature T or B cells [68]. The percentage and absolute number of NK cells in PEC were significantly increased in both infected groups compared to noninfected controls. There was no significant difference in the percentage of NK cells in infected female and male mice (Figure 4A and B) on day 5 post-infection. Surprisingly, the absolute number of NK cells in infected males was higher than in female mice (Figure 4B). The percentage and absolute number of IFNγ+ NK cells at the peritoneum increased (Figure 4C and D), but there was no difference in the percentage and absolute number of IFNγ+ NK cells in female compared to male mice. There was also no difference in the fold increase in either the percentage or the absolute number of NK cells between sexes following infection (Figure 4E). Similar to the fold increase of NK cells, the fold change of the percentage of IFNγ+ NK cell was not different, but the fold change in an absolute number of IFNγ+ NK cells was nearly 2-fold higher in infected males than in infected females (Figure 4F).

**Figure 4.**
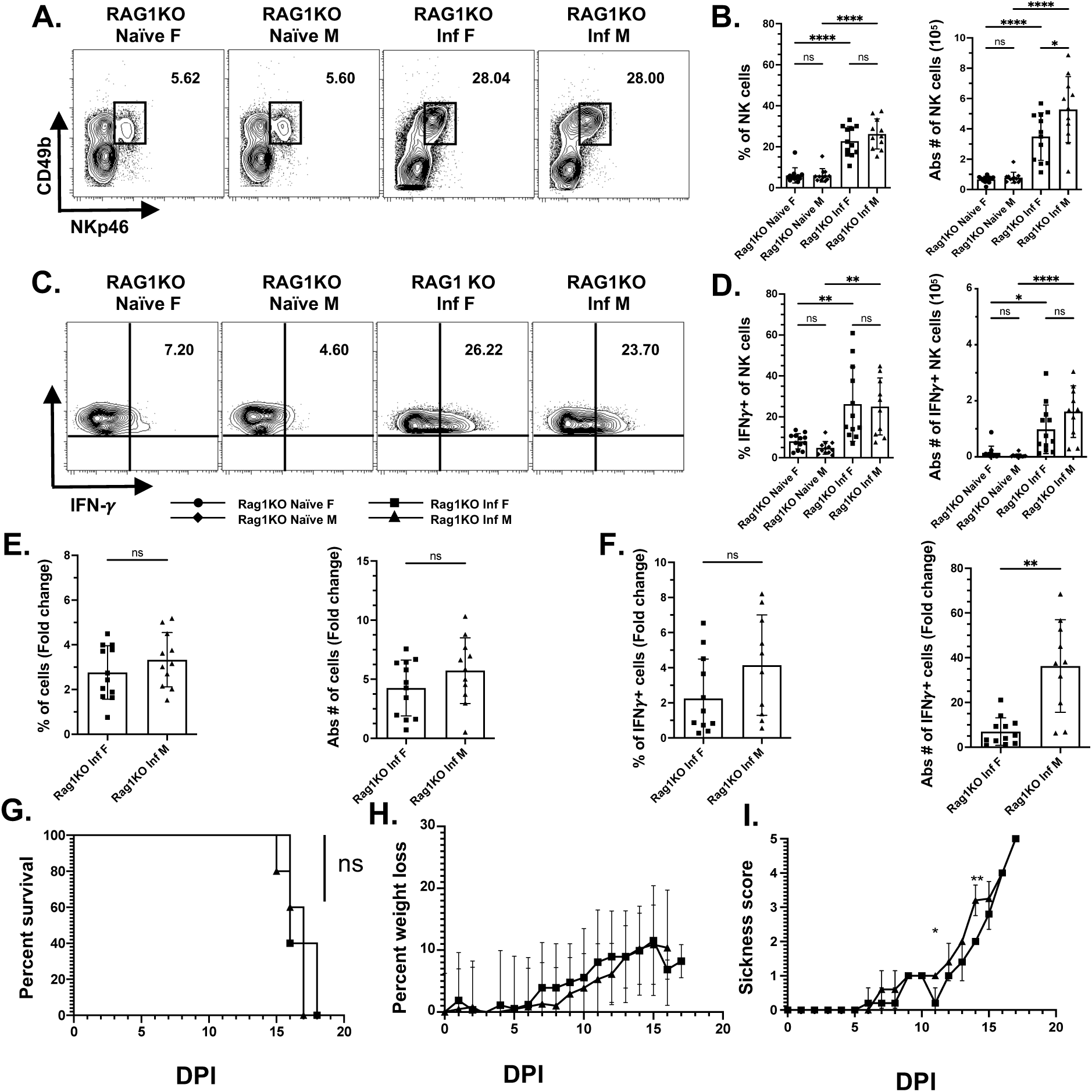
Mature lymphocytes are not involved in sexual dimorphic NK cell response during acute T. gondii infection. WT and RAG1KO female mice were infected intraperitoneally with 20 cysts of *T. gondii* (type II strain, ME49). Five days post-infection, peritoneal cells were harvested and stained for Live/Dead, CD3, NKp46, CD49b, and IFN*γ* to identify NK cells using flow cytometry. Female and male NK cells (Lin-NKp46+CD49b+) and IFN*γ*+NK cells (Lin-NKp46+CD49b+IFN*γ*+) were quantified at the infection site in each group. Representative contour plots are shown for **(A.)** peritoneal NK cells and **(C.)** peritoneal IFN*γ*+ NK cells. The percentage and absolute number of **(B.)** NK cells and **(D.)** IFN*γ*+NK cells in the peritoneum are depicted. Data were pooled from 4 individual experiments. Percentage fold change and absolute number fold change of **(E.)** NK cells and **(F.)** IFN*γ*+NK cells in the peritoneum are presented. RAG1KO female and male mice were infected with 2-3 cysts of *T. gondii* (type II strain, ME49). **(G.)** Percent survival of RAG1KO female and male mice (n=5), **(H.)** percent weight loss, and **(I.)** sickness score (n=10) was evaluated. The log-rank (Mantel-Cox) test was used to assess survival rates. Ordinary one-way ANOVA was employed to analyze the NK cell population. Data are presented as mean ± SD. *p < 0.05, **p < 0.01, ***p < 0.001, ****p < 0.0001.

We then tested how the absence of adaptive lymphocytes results in differential sex-specific survival outcomes in *T. gondii* infection. RAG1KO female and male mice were infected orally with 2-3 cysts of ME49. In the absence of mature lymphocytes or RAG1, female and male mice both succumbed to the low dosage of *T. gondii* infection at the same time, and there were no significant differences detected between the sexes (Figure 4G). Mice started losing weight after day 7 post-infection, with the sickness score following the same pattern (Figure 4H). There was no significant difference across the sexes, except on days 11 and 14 post-infection, when females became sicker than males (Figure 4I). These results combined suggest that mature lymphocytes are not involved in sexual dimorphic NK cell response during acute *T. gondii* infection.

### Group1 ILC Eomes and T-bet do not impact differences in sex-specific survival differences during T. gondii infection

T-box transcriptional factors such as Eomes and T-bet play a crucial role in the development, maturation, and function of group 1 ILCs (NK and ILC1s) [69–72]. *T. gondii* infection can cause NK cells to transform into an ILC1-like phenotype [51]. This phenotype appears to have a negative regulatory role on exhausted CD8+ T cells during persistent infection in an NKp46-dependent manner [27, 57]. Early production of IFNγ by ILC1 and NK cells is dependent upon T-bet during *T. gondii* infection [73]. To further investigate how sex impacts NK cell responses to *T. gondii* infection, we generated group 1 ILC-specific knockdowns of Eomes and T-bet using NKp46iCre transgenic mice [74]. Female and male NKp46iCre^+/−^Eomes^f/f^ and NKp46iCre^+/−^T-bet^f/f^ mice were orally infected with *T. gondii* and monitored for disease outcomes and survival. There were no significant differences in survival when compared to aged-matched NKp46iCre^−/−^Eomes^f/f^ female and male control mice (Supplementary Figure 2A). On day 7 post-infection, both female and male NKp46iCre^+^Eomes^f/f^ showed a significant increase in sickness score, but there was no difference among the sexes except on days 36-37, 52-57, 59-61, 64-69, and 71 males looked sicker than females (Supplementary Figure 2C). Female and male NKp46iCre^+/−^Eomes^f/f^ and NKp46iCre^+/−^T-bet^f/f^ mice followed a similar pattern in weight change (Supplementary Figure 2E and F). However, the percent weight loss in NKp46iCre^+/−^Eomes^f/f^ males tended to be greater than in female mice during the length of the experiments (Supplementary Figure 2G). NKp46iCre^+/−^Eomes^f/f^ mice are considered to be NK cell-deficient mice due to the requirement of the transcription factor Eomes in their development [69, 72, 74–76]. To examine the impact of NK cell loss in female versus male mice during acute *T. gondii* infection, NKp46iCre^+/−^Eomes^f/f^ female and male mice were infected orally (20 cysts ME49) alongside their littermate controls, and parasite burden was measured on day 12 post-infection by real-time PCR. Although there was a difference in parasite burden between NKp46iCre^−/−^Eomes^f/f^ female and male mice, with females showing a higher parasite burden in the mesenteric lymph nodes (MLN) on day 12 post-infection (Supplementary Figure 2I), the loss of NK cells did not correlate with an increased parasite burden in the MLN at this time point. Similarly, no differences in parasite burden were observed across the groups in organs such as the small intestine, spleen, liver, and lungs (data not shown). These results collectively suggest that the transcription factors Eomes and T-bet in group 1 ILC compartment may not explain the sex-specific differences in survival outcomes in *T. gondii* infection.

### IL-12p40 secretion is independent of CD4 co-receptor expression during acute T. gondii infection

NK cell activation during *T. gondii* infection is cytokine dependent, and IL-12p40 is essential for this process [16, 25, 27]. To further investigate a mechanism that explains the difference in NK cell and IFNγ+ NK cell numbers in female CD4 co-receptor KO mice compared to males, we measured the level of IL-12p40 in PEC and spleen. WT and CD4 co-receptor KO female and male mice were infected with 20 cysts of *T. gondii* ME49 i.p. Five days post-infection, PEC and spleen were evaluated for IL-12p40 level by ELISA. As expected, infection increases IL-12p40 levels in the peritoneum as well as in the spleen (Figure 5A and B) in both sexes. However, the absence of the CD4 co-receptor in the system did not significantly reduce IL-12p40 production in either females or males. These data suggest IL-12p40 secretion is independent of CD4 co-receptor expression and not a factor that explains the difference in NK cell responses in female CD4 co-receptor KO mice during acute *T. gondii* infection.

**Figure 5.**
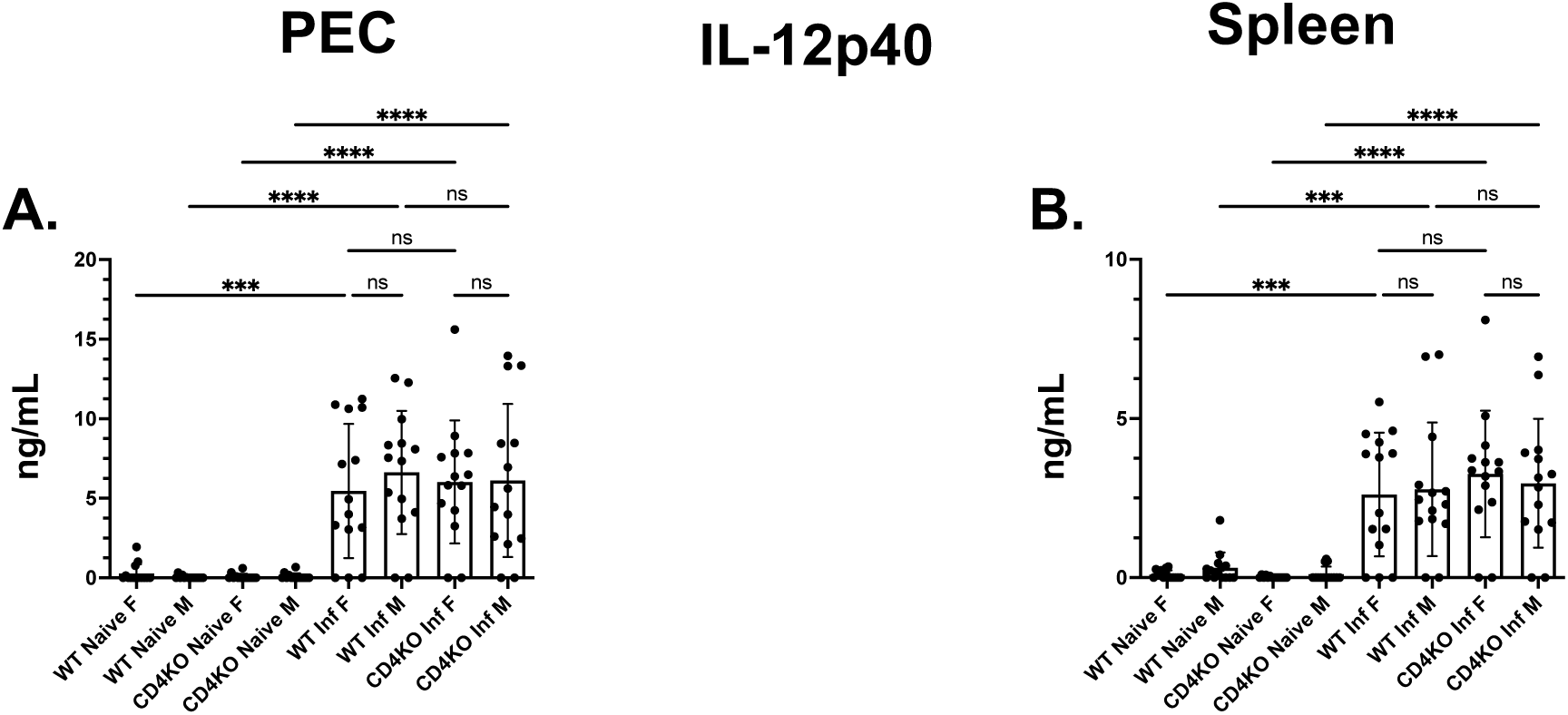
IL-12p40 secretion is independent of CD4 co-receptor expression. WT and CD4KO female and male mice were infected with 20 cysts of *T. gondii* (type II strain, ME49). Five days post-infection, peritoneal cells were harvested and plated using FACS media (2×10^6^ PEC cells and 5×10^6^ splenic cells). The suspended cells were incubated for 24 hours at 37°C with 5% CO_2_. Supernatants were collected, and IL-12p40 ELISA was performed (n=14). Representative graphs of **(A.)** peritoneal and **(B.)** splenic IL-12p40 levels were pooled from 3-4 independent experiments. Ordinary one-way ANOVA was used to evaluate the IL-12 levels in different groups. The data are ± SD *p< 0.05, **p<0.01, ***p<0.001, ****p<0.0001.

## Discussion

Sexual dimorphisms in the immune system exist and have a significant impact on immune responses and the immune protection and pathology they create [77–83]. Understanding how these differences occur is critical for improving the health of female and male patients. Females are more susceptible than males to *T. gondii*, which may result from different immunological responses [45–47]. Female HIV/AIDS patients are also more susceptible to complications with *T. gondii* infection than males [48], suggesting that CD4+ T cells may be a factor in sexually dimorphic immunity. Here, we investigated sexual dimorphism in innate immunity to *T. gondii* infection. Our results demonstrate that CD4 co-receptor is required in a sexually dimorphic manner for optimal NK cell responses during parasite infection in female but not in male mice. Interestingly, NK cell responses were not significantly different between female and male wild-type mice. CD4 co-receptor knock-out (CD4 receptor KO) female mice succumbed to *T. gondii* infection faster than male knockout mice. Despite differences in NK cell responses and survival outcomes in CD4 co-receptor KO female mice, there were no significant differences in NK cell responses when comparing female and male MHCIIKO mice that completely lack CD4+ T cells. Furthermore, no differences in NK cell responses were observed in female RAG1KO compared to male RAG1KO mice. To test how transcription factors important for NK cell development and maturation impact sexually dimorphic NK cell responses, we test how group 1 ILC-specific Eomes and T-bet impact *T. gondii* infection outcomes. Surprisingly, we observed no differences between sexes and no differences between gene knockouts in survival, disease behavior, or parasite burdens. Lastly, we find that IL-12p40 was produced at similar levels in CD4 co-receptor KO female and male mice in PEC and spleen. Taken together, these results suggest that CD4 co-receptor expression is essential for optimal NK cell responses only in female mice, and the difference is not dependent upon transcription factors and cytokines that are required for NK cell responses to infection.

Immune responses to *T. gondii* infection can be different depending upon the sex of the host. The immunological differences between opposite sexes are not well known. Previous studies demonstrate that female BALB/c mice were more prone to *T. gondii* infection than male mice. These differences were attributed to delayed T-cell responses in female mice and lower levels of IFNγ compared to males. In addition to being more susceptible to acute *T. gondii* infection, females (129XB6, B10, and BALB/K strains) also had higher cyst burdens in their brains during chronic infection than males [45]. Other studies in SCID mice demonstrate that males develop more rapid production of IL-12 and IFNγ and as a result, had a lower parasite burden and longer survival times than female SCID mice [47]. Since SCID mice lack B or T cells, the sex-specific differences in infection outcomes could be explained by differences in NK cell responses. However, our data indicate that NK cell responses, including increases in NK cell number and their IFNγ production, are not different between WT female and male mice. Only when the CD4 co-receptor is absent do we detect sex-based differences in NK cell responses. Our work was also performed in a different mouse background, the C57BL/6J background. However, there is confusion about what sex-based differences there were with the initial studies in BALB/c and SCID mice. A recent publication with BALB/c mice repeating what was previously published, but with a new approach using an IVIS whole-body imaging system and luciferase-expressing parasites, demonstrates that females have lower parasite burdens due to developing a more robust immune response during acute infection [46]. However, they have increased morbidity and mortality compared to male mice. We observed female C57BL/6J mice have lower parasite burdens than male mice during acute infection. Despite differing results in previous studies, our results may demonstrate another level of complexity in understanding sexually dimorphic immune responses during *T. gondii* infection. This could be in part due to the genetic background of the animals studied as well as additional factors such as CD4 co-receptor expression in this process

NK cells and adaptive immune cells have a complex relationship where they can regulate each other. NK cells can positively and negatively regulate adaptive immune cells [29, 30, 32, 57, 84–90]. This regulation can occur both during acute and persistent infections, including *T. gondii* infection [57, 91]. CD4+T cells can either directly or indirectly affect NK cell function in different diseases and infections [39–42]. In vitro, upon direct IL-2 stimulation from T cells, human CD56^bright^ NK cells produce IFNγ [37]. In Influenza A (fluA) viral infection, virus-specific T cells from fluA-infected donors’ PBMCs can increase the IFNγ response of CD56^dim/bright^ NK cells through IL-2 [38]. CD4+ T cells also secrete IL-2 in an MHCII-dependent manner to promote IFNγ response from NK cells [39]. Additionally, antigen-specific CD4+ T cells are vital for early NK cell IFNγ secretion in *L. major*-infected mice by directly producing IL-2 and indirectly promoting IL-12 production via CD40/CD40L interactions with DCs [40]. Antigen-specific memory CD4+ T cells are essential for optimum NK cell activation during *P. murina* infection. This results in NK cells upregulating NKG2D and producing IFNγ, granzyme B, and perforin [42]. In a cancer model, CD4+T cells, in conjunction with NK cells, control tumor growth, suggesting that CD4+T cells synergize with NK cells to control tumor cells [92, 93]. B cells can also crosstalk with CXCR5+ NK cells in the simian immunodeficiency virus (SIV) infection model [58]. The interaction between B cells and NK cells is bidirectional. NK cells can positively or negatively regulate B cell activation, function, memory formation, and ag-specific repertoire switching [59–64]. In chronic viral infection, B cells regulate NK cells by upregulating Qa-1b, a ligand for inhibitory NK cell receptors, thereby limiting NK cell activity to promote anti-viral T cell immunity [65]. Despite what is known about CD4+ T cells and their direct and indirect mechanisms of NK cell regulation, we observed that sexually dimorphic NK cell responses and survival outcomes were not apparent in the absence of CD4+ T cells (MHCIIKO). Therefore, the direct or indirect mechanisms by which CD4+ T cells regulate NK cell responses may not be involved in NK cell responses to acute *T. gondii* infection. Additionally, RAG1KO mice that lack not only CD4+ T cells, but also CD8+ T and B cells indicate that sexually dimorphic NK cell responses are independent of the adaptive immune cells.

The influence of recombination-activating gene (RAG) extends beyond adaptive immune cell development and is thought to also contribute to NK cell development, fitness, and functional diversity [94]. Lymphocytes and NK cells are derived from common lymphoid progenitor (CLP), and cytotoxic CD8+T (CLTs) cells share functional, developmental, and educational similarities with NK cells [95]. Unlike T or B cells, NK cells do not undergo VDJ recombination for the receptor rearrangement process. Interestingly, transient expression of RAG proteins and occurrences of unfinished V(D)J recombination have occasionally been observed in a small subset of NK cells during their development, which impacts their function later after development [96–99]. We observed that NK cell and IFNγ+ NK cell numbers increased robustly in RAG1KO mice in both sexes. RAG1KO mice developed a much stronger NK cell response than WT mice (data not shown). This observation is consistent with previous work, which showed that RAG1KO mice have higher numbers of NK cells in the spleen than WT mice [100]. We also observed that *T. gondii* infection of male RAG1KO mice resulted in a higher number of NK cells and a higher fold increase in IFNγ+ NK cells compared to female mice. These results could indicate sex-specific differences in NK cell death, recruitment, or proliferation in the absence of RAG. Another possible explanation for the elevated fold change increase in IFNγ+ NK cells in males compared to females might be due to the sex-specific early requirement of T and B cells in NK cell development and education. Even though *T. gondii* infection of RAG1KO mice resulted in robust NK cell responses, RAG1KO mice still died very early, depicting the importance of adaptive immunity in controlling *T. gondii* infection, and there was no difference in survival between the sexes. Regardless, how the lack of T and B cells and RAG enzymes impacts NK cell development, education, and function is unclear, specifically in a sex-dependent manner.

Eomes and T-bet play a pivotal role in the development, maturation, and function of group 1 innate lymphoid cells (ILCs), including NK cells and ILC1s [69–72, 75, 101–104]. In *T. gondii* infection, these two transcription factors are crucial since they control IFNγ production by ILC1s and NKs during the acute phase of the infection. T-bet-dependent IFNγ secretion by ILC1s and NK cells is critical for maintaining IRF8+ inflammatory cDC1s at infection sites, helping host resistance against *T. gondii* infection [73]. Moreover, *T. gondii* infection induces a transformation of NK cells into cells resembling ILC1 [51]. We investigated how selective tissue-specific knockdown of these two transcription factors in the group 1 ILC compartment (NKp46iCre^+/−^Eomes^f/f^ and NKp46iCre^+/−^T-bet^f/f^) might impact sexually dimorphic resistance to infection. Our results did not show any significant sex-specific differences in survival or disease progression during *T. gondii* infection. These results suggest that other mechanisms may compensate for the lack of transcriptional control by Eomes and T-bet for group1 ILC responses during parasite infection and the absolute requirement of these two transcriptional factors for group1 ILC development, maturation, and function [72, 105, 106]. Like Eomes and T-bet, IL-15 is critical for NK cell development, maturation, proliferation, and function [107–110]. Lack of IL-15 and IL-15 receptor α-chain (IL-15Rα) results in extremely low NK cell numbers in mice [111, 112]. Surprisingly, *T. gondii* infection of IL-15 KO mice that were thought to lack NK cells resulted in robust NK cell responses similar to WT mice [113]. This was later shown to be dependent upon IL-12 [114]. Therefore, another possibility is that knocking down these transcription factors Eomes and T-bet from NKp46+ cells can eliminate NK and ILC1s for a brief period, and upon infection with *T. gondii*, group1 ILCs repopulate and restore their normal function due to inflammatory cytokine milieu present during infection. The growing knowledge that the immunological response to parasites like *T. gondii* is extremely complex, with several layers of regulation and backup mechanisms to maintain host survival, lends even more credence to the idea of redundancy. The complex regulation compensatory networks of the host immune system might explain why deleting critical transcription factors like Eomes and T-bet did not lead to the anticipated detrimental effects on survival or pathogen control. Our findings again emphasize the need for more research into the compensatory mechanisms that prevent infection when essential transcription factors are missing.

The sex-specific requirement of CD4 co-receptor but not CD4 +T cell suggests a role for CD4 co-receptor in the myeloid compartment or on NK cells themselves. It is well established that IL-12 is required for early ILC-1 and NK cell activation in *T. gondii* infection [21, 40, 73, 115], and IL-12 is primarily produced by the myeloid population [14, 116–118]. Upon *T. gondii* infection, pathogen sensing was recognized by myeloid cells via PRRs, like Toll-like receptors (TLRs). These TLR and pathogen antigen interaction triggers the necessary cytokine responses to control the infection [119–124]. CD4 co-receptor can be expressed on different myeloid cells in mice and humans [125–128]. Mouse spleen has a CD4+8-DEC-205^low^ CD11b^high^ DC population [125]. In C57BL/6J mice and humans, macrophages can also be CD4 coreceptor positive [126, 127, 129]. Additionally, human peripheral blood neutrophils can also express CD4 co-receptors [128]. The role of CD4 co-receptor on these myeloid cell populations is not well understood but could contribute to their cytokine production. Our results suggest that CD4 co-receptor did not affect myeloid cell IL-12 production because we did not detect any differences in IL-12p40 production when comparing CD4 co-receptor KO mice with WT mice or in comparing females to males in these different mouse genotypes. Another possibility about the sexually dimorphic NK cell differences in CD4 co-receptor mice could be due to CD4 co-receptor expression on the NK cells themselves. Human NK cells express CD4 co-receptors in blood and lymphoid tissues [130, 131]. These CD4+NK cells isolated from lymphoid tissues show higher expression of CD4 co-receptors than peripheral CD4+NK cells and maintain robust cytotoxicity by producing more IFNγ than CD4− NK cells. Also, ligation of the CD4 co-receptor enhances NK cell’s ability to increase cytokine production and facilitate migration towards IL-16 gradient, highlighting the role of CD4 co-receptor as a chemotactic receptor in innate immunity [131]. This might explain why we noticed fewer NK cells and IFNγ+ NK cells at the site of infection in CD4 co-receptor KO female mice, emphasizing the role of CD4 co-receptors in NK cell migration. Although the cells getting into the site of infection show normal IFNγ responses compared to WT female mice. These observations highlight the complex nature of innate immune signaling in *T. gondii* infection and potential areas for therapeutic targets.

This study investigates sex-specific immune responses to acute *T. gondii* infection, focusing on the role of CD4 co-receptor and NK cell function in both female and male mice. Our findings reveal for the first time that the CD4 co-receptor is crucial for optimal NK cell activity, and remarkably, this requirement is distinctly sex specific. Combined, these findings reveal crucial insights into sex-specific immune regulation during *T. gondii* infection, highlighting the role of CD4 co-receptors in the female’s NK cell function. Our study offers insight into reasons for plausible pathogen vulnerabilities in immunocompromised such as HIV/AIDS populations. This research not only highlights the sexual dimorphic need for CD4 co-receptors, which correlates to poor survival outcomes in female immune-compromised population, but also reveals the intrinsic requirement of CD4 co-receptors for NK cell function during acute *Toxoplasma* infection.

## Acknowledgments

J.P.G. is the principal investigator and corresponding author of this research project. J.P.G. and T.R. comprehended the project. T.R. was responsible for experimental design. T.R., L.B., K.F., H.K.K., and S.K.N. conducted the experiments. T.R. performed data analysis. J.P.G. provided intellectual input and a technical guide. T.R. and J.P.G. drafted the manuscript, with all authors contributing to revisions. J.P.G. was the project supervisor, and J.P.G. secured funding acquisition. We would also like to thank Dr. Stephen L. Denton for his valuable intellectual and technical contributions to the project.

This work was supported by the NIH Wyoming INBRE 2P20GM103432 Pilot project awarded to J.P.G. and the NIH Wyoming INBRE 2P20GM103432 graduate student GA awarded to T.R. This project was also supported by grants NIH R21AI159200 and R21AI161298 awarded to J.P.G.

## Abbreviations List

AIDS: Acquired Immunodeficiency Syndrome
APC: Antigen Presenting Cells
CLP: Common Lymphoid Progenitor
cDC: Conventional Dendritic Cells
DPI: Days Post Infection
DMEM: Dulbecco’s Modified Eagle Medium
FACS: Fluorescence-Activated Cell Sorting
FBS: Fetal Bovine Serum
IACUC: Institutional Animal Care and Use Committee
IFNγ: Interferon-Gamma
IL-15Rα: IL-15 receptor α-chain
ILC: Innate Lymphoid Cells
KO: Knockout
*L. major*: *Leishmania major*
MHC: Major Histocompatibility Comple
MLN: Mesenteric Lymph Nodes
NK: Natural Killer
*P. falciparum*: *Plasmodium falciparum*
*P. murina*: *Pneumocystis murina*
PRR: Pathogen Recognition Receptors
PEC: Peritoneal Exudate Cells
PBS: Phosphate Buffered Saline
PTIC: Protein Transport Inhibitor Cocktail
RAG: Recombination-Activating Gene
RT: Room Temperature
TE: Toxoplasmic Encephalitis
*T. gondii*: *Toxoplasma gondii*
TLR: Toll-Like Receptor
T-bet: T-box transcription factor expressed in T-cells
WT: Wild-Type

