## supplemental figures 1-3 for "CD4 Co-Receptor Regulates Sex-Specific NK Cell Responses to Acute *Toxoplasma gondii* Infection"

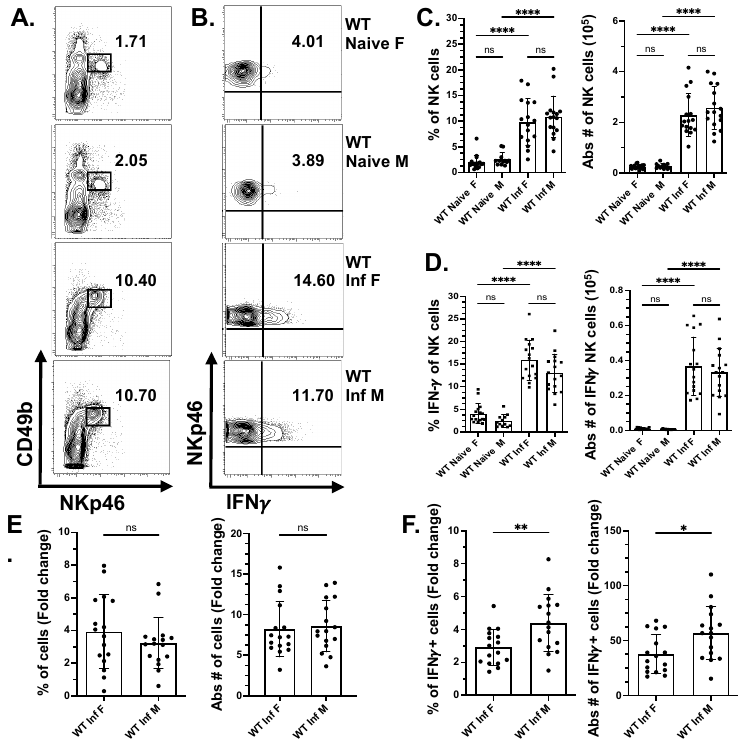

Supplementary Figure 1

*NK cell response is not sexually dimorphic during T. gondii infection when the CD4 co-receptor is expressed.*

WT female and male mice were infected intraperitoneally with 20 cysts of *T. gondii* (type II strain, ME49). Five days post-infection, peritoneal cells were harvested and stained for Live/Dead, CD3, NKp46, CD49b, and IFN𝛾 to identify NK cells using flow cytometry. Female and male NK cells (Lin- NKp46+CD49b+) and IFN𝛾+NK cells (Lin-NKp46+CD49b+IFN𝛾+) were quantified at the infection site in each group. Representative contour plots are shown for **(A.)** peritoneal NK cells and **(B.)** peritoneal IFN𝛾+ NK cells. The percentage and absolute number of **(C.)** NK cells and **(D.)** IFN𝛾+NK cells in the peritoneum are depicted. Percentage fold change and absolute number fold change of **(E.)** NK cells and **(F.)** IFN𝛾+NK cells in the peritoneum are presented. Data were pooled from 4 individual experiments. Ordinary one-way ANOVA was employed to analyze the NK cell population. Data are presented as mean ± SD. *p < 0.05, **p < 0.01, ***p < 0.001, ****p < 0.0001.

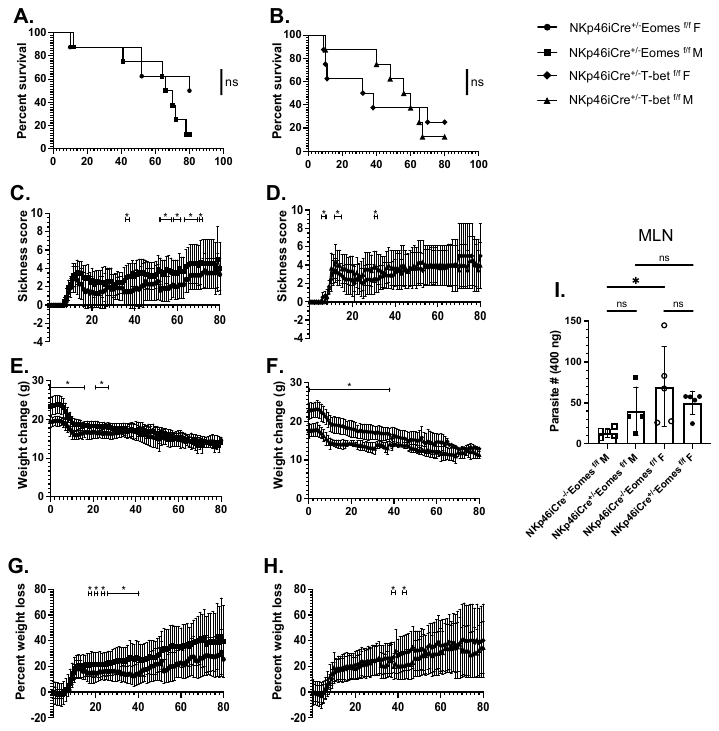

Supplementary Figure 2

*Group1 ILC Eomes and T-bet do not impact differences in sex-specific survival differences during T. gondii infection.*

NKp46iCre^+/-^Eomes ^f^/^f^, NKp46iCre^+/-^T-bet ^f^/^f^ female and male mice were infected orally with 8 cysts of *T. gondii* (type II strain, ME49). **(A.)** Percent survival, **(C.)** sickness score, **(E.)** weight change, and **(G.)** percent weight loss was monitored till day eighty post-infection in NKp46iCre^+/-^Eomes ^f^/^f^ female and male mice (n=8). Similarly, **(B.)** percent survival, **(D.)** sickness score, **(F.)** weight change, and **(G.)** percent weight loss was monitored till day eighty post-infection in NKp46iCre^+/-^T-bet ^f^/^f^ female and male mice (n=8). **(I.)** Graph present *T. gondii* parasite burden of NKp46iCre^+/-^Eomes ^f^/^f^ female versus male mice on day twelve post-infection (oral infection with 20 cysts of ME49, mice n=4-5) in Mesenteric Lymph Node. The log-rank (manel-Cox) test was used to evaluate survival rates. Ordinary one-way ANOVA (Brown-Forsythe and Welch) was used to evaluate the parasite burden. Data are ± SD. *p<0.05, **p<0.01, ***p<0.001, ****p<0.0001.

Supplementary Figure 3

*Gating Strategy for NK Cell and PEC Composition*

C57BL/6 mice were infected intraperitoneally with 20 cysts of *T. gondii* (type II strain, ME49). Five days post-infection, peritoneal cells were harvested and stained for Live/Dead, CD3, NKp46, CD49b, and IFN𝛾 to identify NK cells using flow cytometry. **(A.)** The gating strategy presents a combination of Boolean gates and population gates. Live/Dead- and CD3- populations were gated as lineage-negative populations. Population gates include NK cells (Live/ Dead-, CD3-, NKp46+, CD49b+) and IFN𝛾+NK cells (Live/ Dead-, CD3-, NKp46+, CD49b+, IFN𝛾).
